# The unique function of Runx1 in skeletal muscle differentiation and regeneration is mediated by an ETS interaction domain

**DOI:** 10.1101/2023.11.21.568117

**Authors:** Meng Yu, Konrad Thorner, Sreeja Parameswaran, Wei Wei, Chuyue Yu, Xinhua Lin, Raphael Kopan, Matthew R. Hass

**Affiliations:** State Key Laboratory of Genetic Engineering, School of Life Sciences, Greater Bay Area Institute of Precision Medicine (Guangzhou), Zhongshan Hospital, Fudan University, Shanghai 200438, China; Division of Developmental Biology, Department of Pediatrics, University of Cincinnati College of Medicine and Cincinnati Children’s Hospital Medical Center, Cincinnati, Ohio, USA; Division of Human Genetics, Department of Pediatrics, University of Cincinnati College of Medicine and Cincinnati Children’s Hospital Medical Center, Cincinnati, Ohio, USA

## Abstract

The conserved Runt-related (RUNX) transcription factor family are well-known master regulators of developmental and regenerative processes. *Runx1* and *Runx2* are both expressed in satellite cells (SC) and skeletal myotubes. Conditional deletion of *Runx1* in adult SC negatively impacted self-renewal and impaired skeletal muscle maintenance. *Runx1-* deficient SC retain *Runx2* expression but cannot support muscle regeneration in response to injury. To determine the unique molecular functions of Runx1 that cannot be compensated by Runx2 we deleted *Runx1* in C2C12 that retain *Runx2* expression and established that myoblasts differentiation was blocked *in vitro* due in part to ectopic expression of *Mef2c,* a target repressed by *Runx1*. Structure-function analysis demonstrated that the Ets-interacting MID/EID region of Runx1, absent from Runx2, is critical to regulating myoblasts proliferation, differentiation, and fusion. Analysis of in-house and published ChIP-seq datasets from *Runx1* (T-cells, muscle) versus *Runx2* (preosteoblasts) dependent tissue identified enrichment for a Ets:Runx composite site in *Runx1*-dependent tissues. Comparing ATACseq datasets from WT and Runx1KO C2C12 cells showed that the Ets:Runx composite motif was enriched in peaks open exclusively in WT cells compared to peaks unique to Runx1KO cells. Thus, engagement of a set of targets by the RUNX1/ETS complex define the non-redundant functions of *Runx1*.

## Introduction

The Runt-related transcription factors (RUNX1, RUNX2, and RUNX3) play essential roles in diverse tissues regulating homeostasis, cell proliferation, lineage specification, and cell fate determination. Runx1 and Runx2 proteins bind to the same DNA site through a highly conserved (>95%) RUNT DNA binding domain at N-terminus, and to the CBFβ transcription factor through the proline-rich (PY) motif. RUNX family members act redundantly in many developmental and disease processes. For example, loss of the pro-oncogenic activity of RUNX1 in acute myeloid leukemia (AML) can be compensated by elevated expression of *RUNX2* and *RUNX3*. Only simultaneous loss of all RUNX activity by reagents designed to bind consensus RUNX-binding sequences or by shRNA lentiviruses targeting all three *RUNX* orthologs provided the desired anti-AML effect (Morita et al., 2017). By contrast, *Runx1* has non redundant activity in T-cell (Taniuchi et al., 2002; Wong et al., 2011) and engages specific targets in mK4 cells (Hass et al., 2021), whereas *Runx2* regulates osteogenesis.

To explore the basis for non-redundancy, we selected to study skeletal muscle. Skeletal muscles account for ∼40%-50% of adult human body weight and are comprised of multinucleated syncytium formed during development by fusion of mononucleated progenitor cells. Muscles have a remarkable capacity for repair and regeneration in response to exercise, injury, aging, or disease, driven by a quiescent resident stem cell population called satellite cells (SC). In response to injury satellite cells are activated, re-enter the cell cycle, and divide asymmetrically to replenish the pool of quiescent stem cells and produce a transiently amplifying population of committed progenitors or myoblasts. The myoblasts expand, then differentiate and fuse with the existing fibers to regenerate a functioning muscle (Brack and Rando, 2012; Lepper et al., 2011; Pawlikowski et al., 2015). Quiescent satellite cells require the paired box 7 (*Pax7*) protein (von Maltzahn et al., 2013), transition to myoblasts is controlled by the sequential expression of myogenic regulatory factors (MRFs), a family of basic helix-loop-helix transcription factors that includes *Myf5*, *MyoD*, followed by *myogenin* and *MRF4* (Penn et al., 2004). The MRFs collaborate with Mef2 family members to drive expression of structural and metabolic genes required in mature muscle fibers (Blais et al., 2005; Braun and Gautel, 2011; Molkentin and Olson, 1996; Potthoff and Olson, 2007).

Genetic analyses established that *Runx1* is activated in injured skeletal muscle and is essential for regeneration: Genome-wide analysis of Runx1-occupied regions coupled with gene-expression analyses revealed enrichment for the RUNX, MyoD and AP-1/c-Jun motifs in primary myoblast (Umansky et al., 2015), and identified key targets including Mef2c and Cdkn1c (p57). Moreover, loss of *Runx1* exacerbates the muscle wasting phenotype of MDX mice leading to early lethality (Umansky et al., 2015). However, the role of Runx1 in maintenance of muscle satellite cells was not analyzed, neither was possible redundancy (or lack thereof) between Runx1 and Runx2 in muscle regeneration.

In this study, we investigated the role of Runx1 protein in muscle homeostasis. We confirmed that conditional deletion of *Runx1* in skeletal muscle satellite cells severely impaired regeneration of injured skeletal muscle and observed that muscle maintenance failed in uninjured muscle due to progressive loss of satellite cells. To address redundancy, we used CRISPR to remove *Runx1* from C2C12 myoblasts, which retain expression of *Runx2*. Loss of *Runx1* led to many transcriptional changes in growth media. In differentiation media, *Runx1*-deficient C2C12 cells failed to differentiate and did not fuse into myotubes. Mef2C was the highest induced gene in Runx1KO C2C12 cells, and over-expression of Mef2c in otherwise wild-type C2C12 cells was sufficient to mimic the fusion defect seen in *Runx1*KO cells. To investigate the molecular basis for the functional differences between Runx1 and Runx2, we generated Runx1/2 chimeras and tested their ability to rescue differentiation of *Runx1*-Null C2C12 myoblasts. Using this assay, we demonstrated that the substitution of the Runx2 MID domain (MID2) with that of Runx1 (MID1) enabled Runx2/1 chimera proteins to rescue *Runx1*-null myoblast proliferation, differentiation and fusion; and conversely, replacement of MID1 with MID2 in Runx1 abolished its ability to rescue. The MID1 domain contains an ETS-interaction helix absent from MID2 suggesting that unique Runx1 functions are Ets-dependent. Co-immunoprecipitation experiments confirmed that in C2C12 cells Runx1 (but not Runx2) can form a complex with the Ets factor Etv4. Finally, analysis of C2C12 ATAC-seq data as well as published ChIP-seq datasets from *Runx1-* or *Runx2*-dependent tissues revealed significant enrichment for a composite Ets:Runx site specifically in *Runx1*-dependent tissues. Altogether, our findings identify Runx1 and Etv4 form a complex regulating Mef2 abundance, satellite cell maintenance, and skeletal muscle regeneration.

## Result

### Deletion of Runx1 in satellite cells impairs skeletal muscle regeneration

Previously, Runx1 function was examined in mice engineered to conditionally remove *Runx1* (*Runx1^fl/fl^*) in myoblasts the presence of Cre recombinase expressed under the control of *Myf5* (Umansky et al., 2015). To investigate *Runx1* function in adult skeletal muscle satellite cells (SCs), we bred *Runx1^fl/fl^* dams with sire carrying a tamoxifen-regulated satellite cell (SC) -specific Cre recombinase (*Pax7^+/CreERT2^: Runx1^+/fl^)* transgene. Following tamoxifen administration, deletion of *Runx1* will occur in both quiescent and activated SCs, where *Pax7* is expressed. The *Pax7^+/CreERT2^*; *Runx1^fl/fl^* (thereafter referred to as *Runx1*cKO) and control *Pax7^+/CreERT2^*; *Runx1^+/fl^* littermates (thereafter referred to as controls) were used for subsequent analyses (Fig. 1A). Without tamoxifen, no differences were observed between *Runx1cKO* and control over their entire lifespan. To confirm that this strategy impacted regeneration after injury in vivo, we administered tamoxifen daily for 5 days to 6-week-old littermates to induce genetic deletion of *Runx1* in SCs, followed by cardiotoxin (CTX) injection into the Tibialis anterior (TA) muscle. We harvested the injured and contralateral control muscles at D5, D7, D12, and D30 post injection (Fig. 1B), and imaged the dissected TA muscles to compare the size (Fig. 1C) and weight (Fig. 1D). Staining for Runx1 and Pax7 in the control and *Runx1*cKO section confirmed Tamoxifen-induced *Runx1* deletion in SCs (Supplemental Fig. S1A): whereas most (∼98%) of Pax7 positive cells in control muscles express Runx1 protein, very few Pax7 positive cells expressed Runx1 in *Runx1* KO muscles (Fig. 1E).

**Figure 1:**
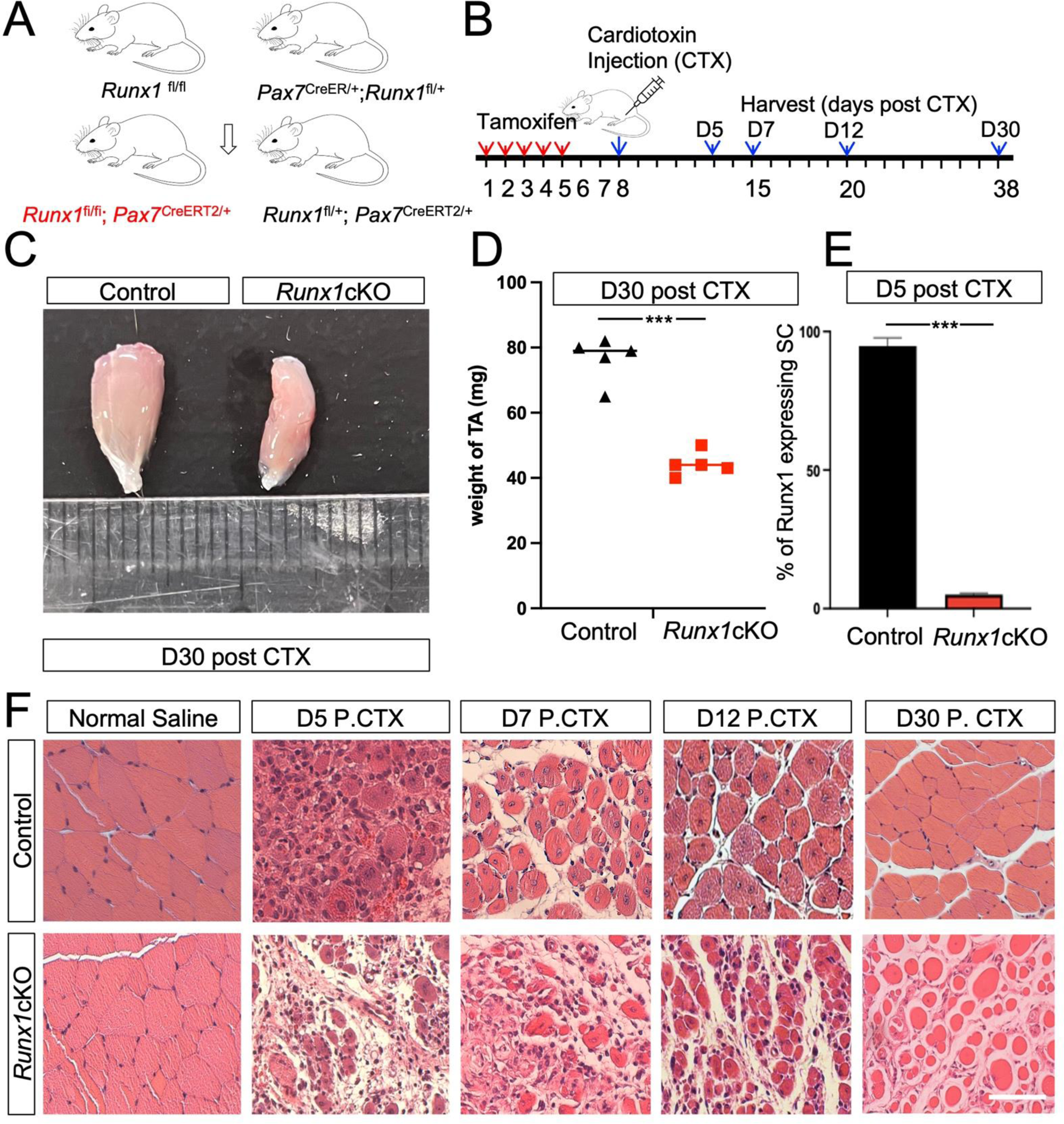
Deletion of *Runx1* in SCs impairs regeneration of skeletal muscle. (A) Breeding scheme to obtain control and Runx1cKO mice (red text). (B) Schematic outline of the strategy for tamoxifen and cardiotoxin administration. See text for detail. (C) Representative image of dissected Tibialis anterior (TA) muscle from control and Runx1cKO mice at D30 post-injury. (D) Measurement and quantification of TA weight in control and Runx1cKO mice (n=5). (E) Quantification of percentage of Runx1-positive cells in Pax7-expressing cells per area from injured TA muscle cross-sections of control and Runx1cKO mice at 5 days post-injury. (F) Cross sections of TA muscles from control and Runx1cKO mice analyzed by H&E staining on days 5, 7, 12, and 30 after saline or CTX injection. Scale bars: 80 μm. Data are represented as mean ± SD. **p<0.01 (Student’s t-test).

Next, we analyzed the regeneration capacity of the injured muscle by histology. Hematoxylin and eosin (H&E) stained sections of control or *Runx1*cKO TA muscle were of indistinguishable size and myofiber architecture in the absence of injury. CTX induced extensive muscle damage and inflammatory infiltrate in both control and *Runx1*cKO muscle after injury. Whereas control mice began to recover by D7 and completed recovery by D30 post-injury (Fig. 1F), *Runx1*cKO muscles still contained degenerating myofibers, fibrotic tissues, and inflammatory cells on D7-post-injury. Very few small regenerating fibers were seen in *Runx1*cKO TA muscle, perhaps arising from SC that failed or had delayed deletion of *Runx1*. Muscle tissue was not reconstituted by D60 in Runx1cKO mice (not shown).

### Runx1 is essential for SC maintenance, proliferation and differentiation

To investigate the function of Runx1 in SC proliferation during development stage, tamoxifen injections were given to pregnant females on day 12.5 of pregnancy (E12.5), which can specifically delete the Runx1 gene in SC during the development of offspring. At the 2^nd^ day after birth (P2), the average body length (L) of Runx1cKO was significantly shorter than controls. The average body weight (W) was ∼62% of that of the control group (Fig. 2B). By P30 the difference between Runx1cKO mice and control was more pronounced (Fig. 2A). These results suggested that Runx1 acted in quiescent or self-renewing SC.

**Figure 2:**
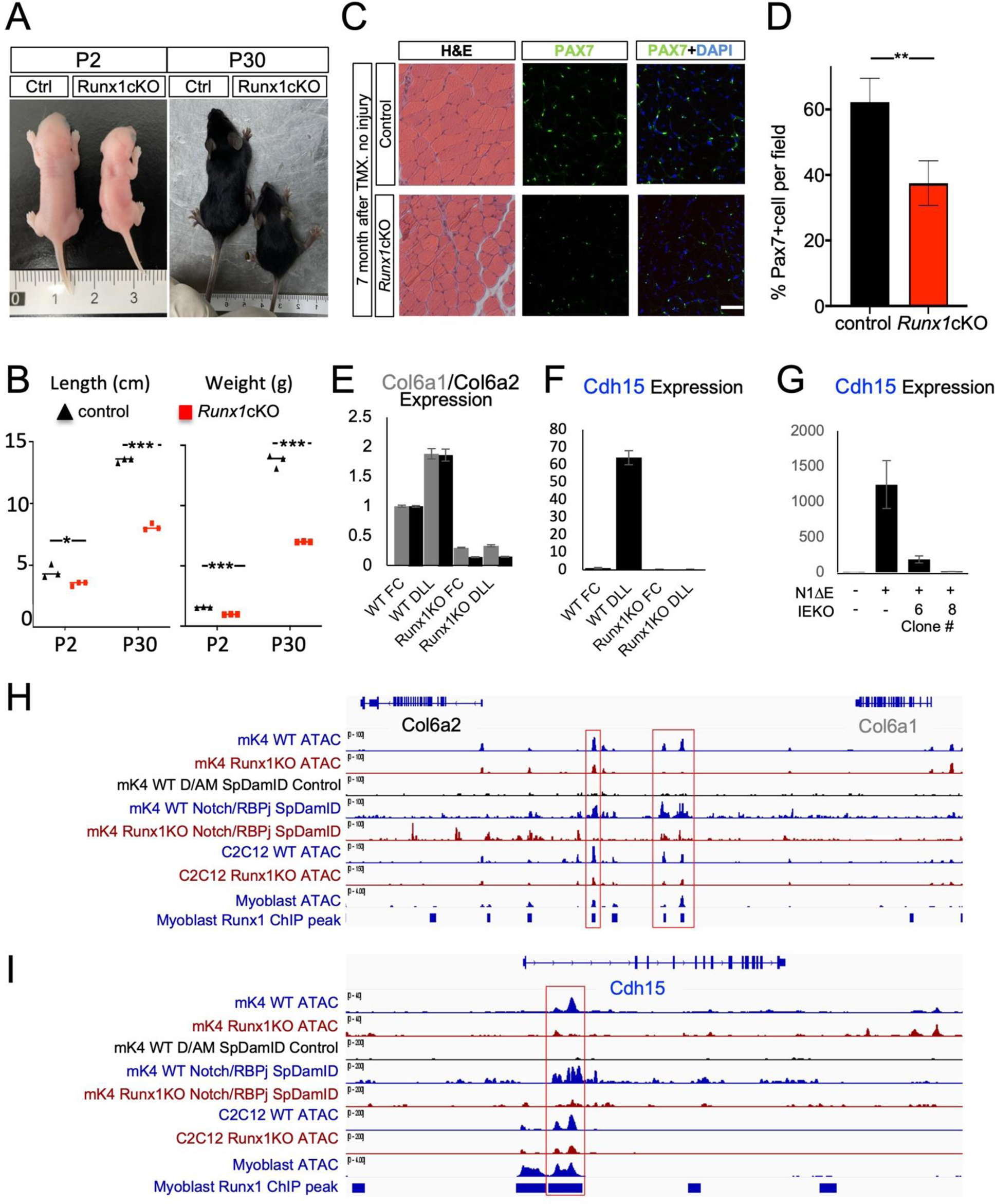
*Runx1* is essential for SC proliferation and maintenance. (A) Representative image of newborn (P2) and weaned (P30) mice exposed to Tamoxifen on day 12.5 of pregnancy (E12.5). (B) Average body length (L) and weight (W) at P2 and P30 for each genotype. Quantification of Body lengths (L) in control and Runx1cKO (n=3). Quantification of body weight (W) in control and Runx1cKO mice (n=3). (C) TA muscle cross-sections at 7 months after tamoxifen administration stained with H&E or Pax7 antibodies (green) and Hoechst (blue) (D) Quantification of Pax7-expressing cells number per area from randomized fields presented as mean ± SD. N=3 for each group. **p<0.01, ***p<0.001 (Student’s t-test). Scale bars: 80 μm. (E and F Bar graphs showing the Runx1-dependent Notch induced expression changes of *Col6a2*, *Col6a1*, and *Cdh15* in control versus Runx1KO mK4 cells exposed to FC or DLL1 ligand for 4 hours. (G) Bar graph displaying that *Cdh15* expression induction by an activated Notch construct (N1ΔE) in cells in which CRISPR Cas9 was used to delete the open intronic region bound by Notch and Runx1 in mK4 and myoblasts. (H and I) Genomic snapshots of the Notch transcriptional targets *Col6a2*, *Col6a1*, and *Cdh15* showing the normalized signal from ATAC-seq from WT (blue) or Runx1KO (red) mK4 cells; Notch complex binding detected by SpDamID in WT (blue) or Runx1KO (red) mK4 cells; ATAC-seq from WT (blue) or Runx1KO C2C12 (red), and ATAC-seq and Runx1 ChIP from primary myoblasts. Red boxes highlight Runx1-bound accessible genomic regions that are reduced or gone in Runx1KO cells.

To investigate the effect of Runx1 deletion on SC cell maintenance we removed *Runx1* from quiescent SC cells by administration of tamoxifen daily for 5 days to 8-week-old control and Runx1cKO littermates. After 7 months the mice were euthanized, and the TA muscle was harvested for analysis. H&E staining showed gaps in fiber packing and smaller cross-sections (Fig. 2C). Pax7 immunofluorescence staining revealed a reduced number of Pax7-expressing SC in Runx1cKO muscles: control mice had an average of 62.25% Pax7-positive SC per field of view, whereas Runx1cKO mice had only 37.5% (Fig. 2D). This result suggests that in uninjured muscle *Runx1* acts to maintain quiescent SC.

Quiescent SC maintenance depends in part on the Notch signaling pathway (Bi et al., 2016; Conboy et al., 2003; Fujimaki et al., 2018; Gioftsidi et al., 2022). Previous studies have shown that Runx1 can affect Notch signaling in T cells (Choi et al., 2017; Wang et al., 2014). We noticed that Runx1 deficient mouse kidney metanephric mesenchyme cell line, mK4 (Hass et al., 2021), shared many open chromatin regions with C2C12 cells suggesting that these cells utilize similar regulatory schemes. During a study on the role of Runx1 in the regulation of Notch signaling we noticed that Notch targets important to SC maintenance, namely the SC marker *Cdh15* and the extracellular matrix proteins *Col6a1* and *Col6a2* (Baghdadi et al., 2018) are robustly of expressed upon exposure of mK4 cells to the Notch DLL1, and that this Notch-dependent induction was abrogated in *Runx1*KO mK4 cells (Fig. 2E and F). We performed ATAC-seq in *Runx1*+ and *Runx1*KO mK4 cells and observed loss of open chromatin in *Runx1*KO cells near these genes (Fig. 2H and I). Furthermore, SpDamID (Hass et al., 2015) demonstrated Notch/RBPj complex binding to these regions in *Runx1*+ but not in *Runx1*KO mK4 cells. These same regions are accessible in myoblasts and recovered by Runx1 in ChIP-seq data from myoblasts (Umansky et al., 2015; Zhang et al., 2020) suggesting that Runx1 is required to maintain accessibility at Notch-responsive enhancers regulating the expression of these targets. To test the assumption that these regions are involved in Notch-dependent regulation, we used CRISPR to delete the *Runx1-*dependent ATAC peak bound by Notch/RBPj in the *Cdh15* intronic region shared in mK4 cells and myoblasts (Supplemental Fig. S2). Deleting the accessible region abolished Notch responsiveness of *Cdh15* in mK4 cells (Fig. 2G). These results support the notion that Runx1 functions in SC maintenance in part by enabling Notch-dependent expression of SC maintenance genes; and in its absence, Runx2 fails to maintain the accessibility of these region.

### Loss of Runx1 impairs C2C12 cell differentiation via dysregulation of Mef2c expression

To further investigate the molecular mechanisms underlying the role of Runx1 in myogenesis we used C2C12 cells, which express both *Runx1* and *Runx2*. We generated *Runx1*KO C2C12 cell lines with CRISPR-Cas9 and guide RNAs targeting exon 3 that encodes part of the DNA binding domain. Western Blot (WB) analyses showed that there was no change in Runx2 expression level in cells lacking Runx1 protein (Fig. 3A). The growth of *Runx1*KO cells in GM was similar to WT cells (Fig. 3B). To differentiate the cells, we switched them from growth medium (GM; Ham’s F10 with 20% FBS, 10ng/mL bFGF and 1% Penicillin/streptomycin (Pen/Strep)) to differentiation medium (DM; DMEM with 2% horse serum and 1% Pen/Strep) when cell density reached ∼ 70% confluence. We then compared their ability to differentiate into myotubes *in vitro*. After ten days in DM, robust differentiation into multinucleated myotubes was observed in controls. By contrast, most of the *Runx1*KO cells failed to fuse (Fig. 3B). 88.67% of WT cells contain 3 or more nuclei, in contrast to only 6.01% in *Runx1*KO cultures (Fig. 3C). These results show that *Runx2* cannot perform the differentiation-promoting functions of *Runx1*.

**Figure 3:**
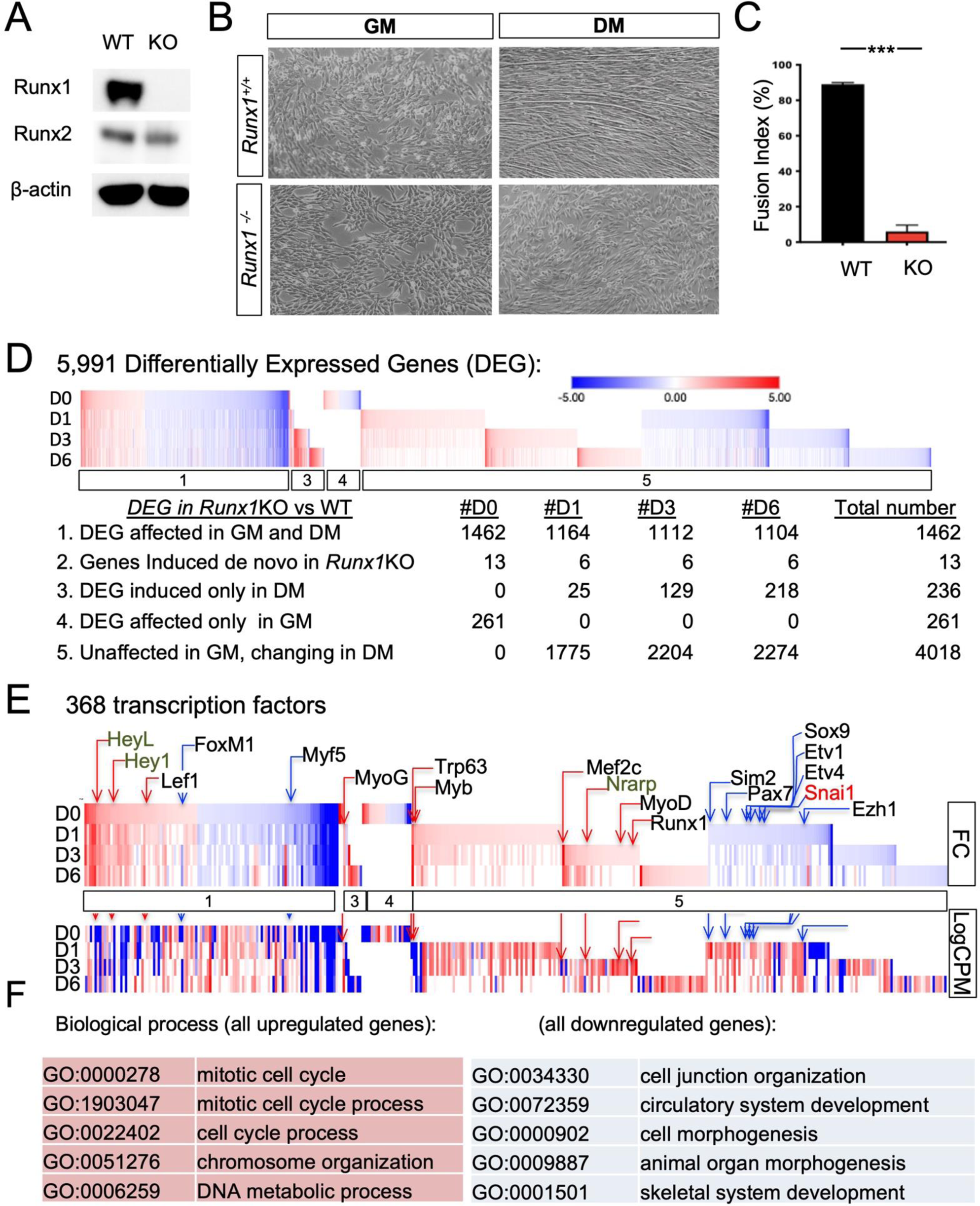
Loss of *Runx1* impairs C2C12 cell differentiation. (A) Western blot analysis of Runx1and Runx2 in WT and Runx1KO C2C12 relative to β-actin. (B) Representative phase contrast microscopy images of WT and Runx1KO C2C12 cells in growth medium (GM) or after 6 d in differentiation medium (DM). (C) Quantification of fusion index of WT and Runx1KO C2C12 cells after 6 days in differentiation medium. ***p<0.001. Data present as means ± SEM. n=3 for each group. (D) Differential gene expression analysis reveal differences in GM (D0) and DM (D1,2,6). The affected genes in each group/day are displayed here and in Supplemental Table S1. (E) Heat maps of the changes in 368 transcription factors: above the clusters, relative Log2 fold change in each column is shown. Below the clusters, the relative abundance (LogCPM) is shown. Members of the Notch pathway, Ets, MRF gene families are indicated above their relative position. (F) GO terms of the up- and downregulated genes in Runx1KO. Details in Supplemental Table S1.

We next analyzed the transcriptome of C2C12 and *Runx1*KO C2C12 cells at GM (D0), and at D1, D3 and D6 in DM (Fig. 3D, Supplemental Table S1, and https://www.ncbi.nlm.nih.gov/geo/query/acc.cgi?acc=GSE248045). 5991 differentially expressed genes passed our cutoff filters (expression >1.7CPM in at least on replicate, fold change (FC) >1.5, pValue <0.01 and FDR <0.05, Table S1). Of these, 1463 genes were differentially expressed in GM (D0; cluster 1 in Table S1 and Fig. 3D), reflecting *Runx1*-dependent changes in myoblasts. 4018 genes that were not Runx1 dependent on D0, increased (1978) or decreased (2040) in expression starting after 24 hours in DM in (D1, Cluster 5, Fig 3D and Supplemental Table S1). Of these, 368 are transcription factors (Fig. 3E and Table S1, column AQ). Notably, the most upregulated transcription factor on D3 was *Mef2c*, which was also upregulated in Runx1-deficient primary myoblasts (Umansky et al., 2015). Other upregulated factors included *Nrarp*, a universal target of Notch signaling, *MyoD*, *Trp63* (promotes differentiation) and *Myb* (promotes proliferation). Downregulated genes included *Pax7*, the Ets factors *Etv1* and *Etv4,* and the repressor *Snai1*. Toppfun analysis of the 1978 upregulated genes revealed enrichment for cell cycle Go-term, whereas the 2040 downregulated genes were enriched for general cellular processes (Fig 3F, Tables S1, Tabs 3, 4). 236 *Runx1*KO C2C12 differentially expressed genes were specific to DM, most were upregulated, and all were enriched for muscle-specific terms (Fig. 3D, cluster 3 in Table S1). The same was true for 1323 downregulated DEG regardless of expression status in GM. Contrary to the observation in primary myoblasts, we do not see strong elevation in Cyclin-dependent kinase inhibitor genes (Supplemental Fig. S3).

To analyze the expression of the MRF *Myogenin, Mef2c, MyoD*, and the differentiation marker *MHC*, mRNA was isolated from cells in GM or after 2, 3 and 6 days in DM. qRT-PCR analysis revealed that *MHC* mRNA levels decreased 80-fold after six days in differentiation medium, consistent with the observed defect in differentiation (Fig. 4A, B). Whereas the mRNA levels of *MyoD* and *myogenin* were not significantly changed, *Mef2c* mRNA levels dramatically increased relative to the WT levels in *Runx1*KO cells after switching to the differentiation medium (Fig. 4A). *Mef2c* mRNA continued to accumulate every day in DM media, exceeding 1000-fold the levels seen in differentiating C2C12 cells (Fig. 4A). Thus, in most respects C2C12 mirror the defect seen *in vivo*, enabling further analysis of the mechanisms involved.

**Figure 4:**
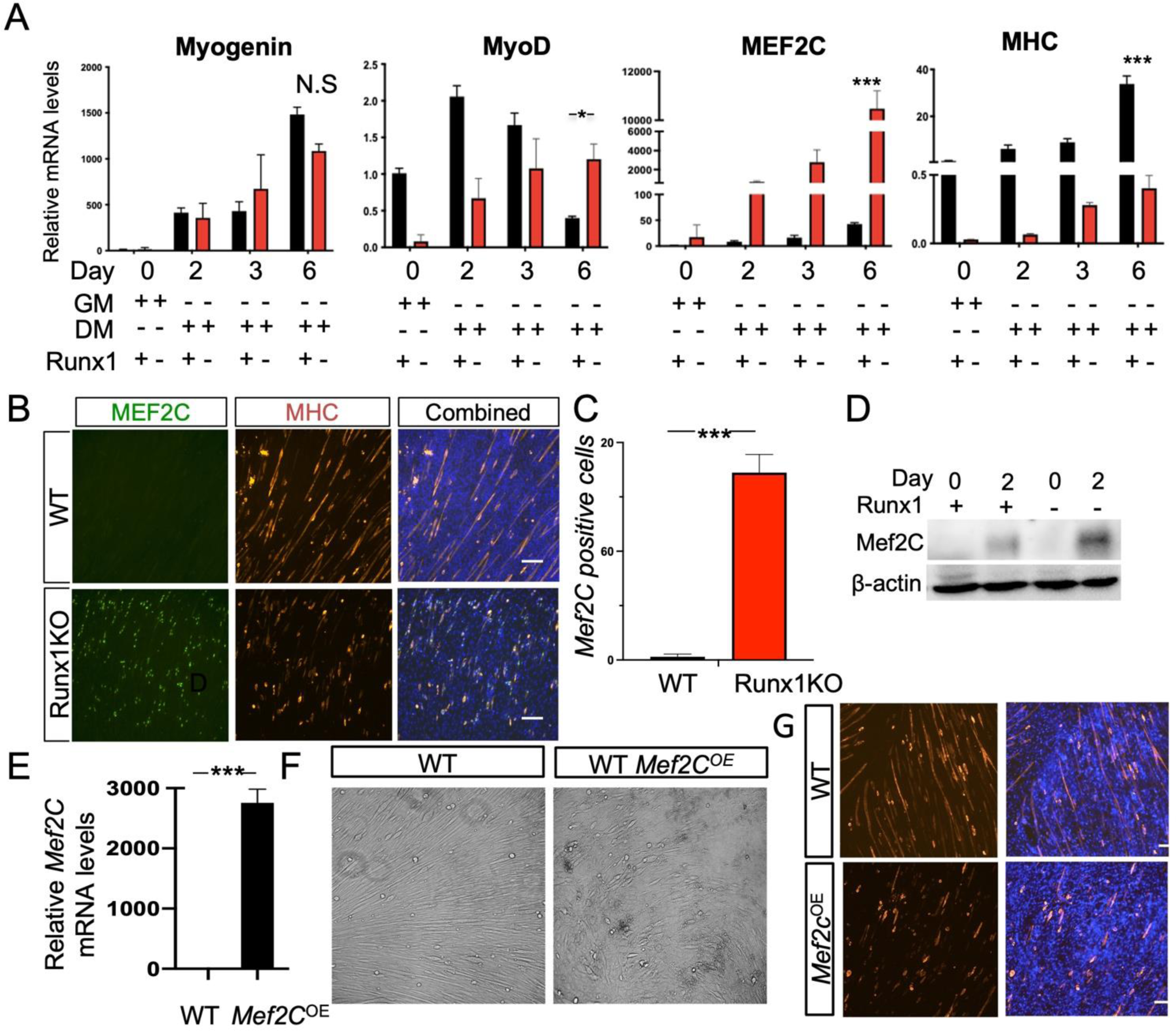
*Runx1* loss blocks C2C12 differentiation via Mef2c. (A) qRT-PCR analysis of Myogenin, MyoD, Mef2c and Myosin Heavy chain (MHC) expression in WT and Runx1KO C2C12 cells on D0, 2, 3, and 6. (B) Immunostaining for Mef2c and MHC WT and Runx1KO C2C12 cells cultured in DM for 6 days. (C) Quantification of the number of mef2c positive cells in (B). (D) Western blot analysis of mef2c expression in WT and Runx1KO cells after 2 days of differentiation. (E-G) C2C12 cells were transfected with negative control or mef2c overexpression plasmid (mef2c-OE) for 24 h in the growth medium and then cultured in the differentiation medium for 6 days. (E) mRNA expression of mef2c in WT and mef2c-OE C2C12 cells. (F) Representative phase contrast microscopy images of WT and mef2c-OE C2C12 cells in differentiation medium for 6 days. (G) Immunofluorescence staining of MHC (red) and DNA (Hoechst, blue) of cells in (F). Data presented as means ± SEM. n=4 for each group. ***p<0.001. N.S., not significant (Scale bars: 80 μm.).

Since mRNA changes are not always concordant with protein levels, we examined Mef2c level through immunofluorescence staining in MHC-positive cells after 6-days in DM. Multinucleated myotubes were evident in control cells but not in *Runx1*KO (Fig. 4B). We counted the number of Mef2c positive cells in both WT and *Runx1*KO cells after differentiation as detailed in methods. The number of strongly positive Mef2c nuclei was significantly higher in *Runx1*KO cells (Fig. 4C). Western blot analysis confirmed the accumulation of Mef2c *Runx1*KO C2C12 grown in DM (Fig. 4D), but to a clearly lesser degree than the mRNA.

Loss of Mef2c impairs muscle differentiation(Liu et al., 2014), but it is unclear of elevated Mef2C levels could do the same. To mimic the effect of losing *Runx1* during differentiation, we overexpressed Mef2c in C2C12 cells (see methods for detail). *Mef2c* mRNA levels were greatly increased in transfected C2C12 (Mef2c^OE^) cells after transfection (Fig. 4E) to a degree similar to that seen in *Runx1*KO cells. Both phase contrast images (Fig. 4F) and immunostaining (Fig. 4G) show that overexpressed Mef2c can block fusion of otherwise WT C2C12. It suggests that the differentiation-preserving function of Runx1 in myoblasts is in part to regulate appropriate expression of *Mef2c*. Clearly, additional activities are likely to be involved in maintenance and proliferation of SC.

### Runx1-specific functions are encoded in the MID domain region

Runx2 is present in C2C12 *Runx1*KO cells and yet it cannot prevent the defect in myogenic differentiation. We first attempted to rescue *Runx1*KO C2C12 cells by overexpressing *Runx1* and/or *Runx2.* Transfecting 200ng of *Runx1* expressing pCDNA3.1 plasmid into *Runx1*KO cells restored differentiation in DM but transfecting 200ng of *Runx2* expressing pCDNA3.1 plasmid did not (Fig. 5A). Fusion index calculation (Fig. 5B) and cell length measurements (Fig. 5C) identified no significant difference between *Runx1*-rescued *Runx1*KO C2C12 cells and WT cells.

**Figure 5:**
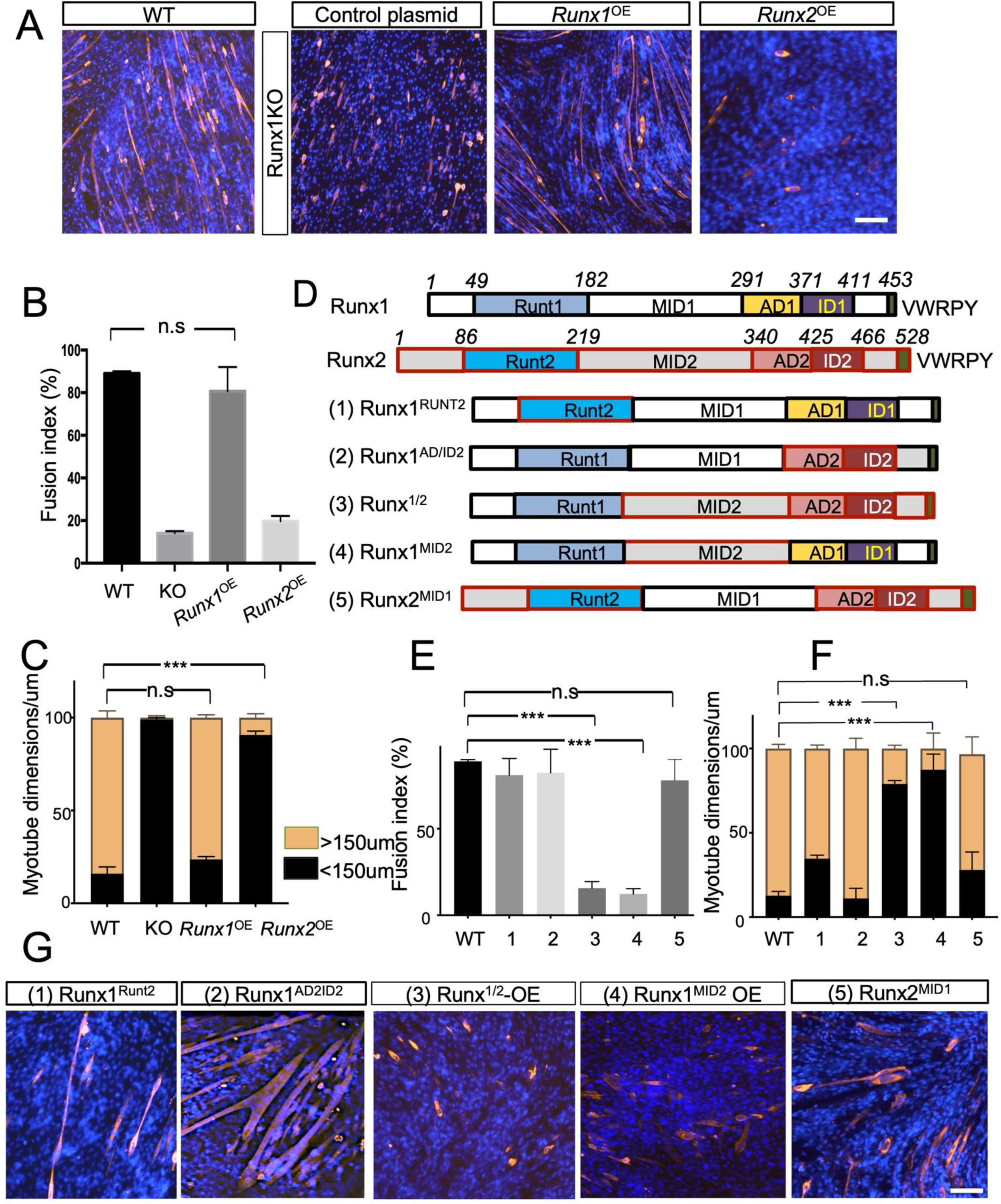
Runx1-specific functions are encoded in the MID domain region. (A) Runx1KO C2C12 cells were transfected with 200ng *Runx1* or *Runx2* plasmids. After 24 hr, cells were induced to differentiate for 6 days followed by immunofluorescence staining for MHC (red) and DNA (Hoechst, blue). (B) Quantification of fusion index and of (C) MHC+ cell lengths in experiments shown in A. (D) Schematics of Runx1 and Runx2 chimera proteins. (E) Quantification of fusion index (F) and MHC+ cell lengths in experiments shown in G; (G) Runx1KO C2C12 cells were transfected with 200ng of five different chimera plasmids numbered in D. 24 h for after transfection, cells were induced to differentiate for 6 days followed by immunofluorescence staining for MHC (red) and DNA (Hoechst, Blue). Data presented as means ± SEM. n=3 for each group. ***p<0.001. n.s., not significant (Scale bars: 150μm.).

The ability of transfected *Runx1* to rescue differentiation enabled structure-function analyses aimed at identifying the domain (s) supporting myogenesis unique to Runx1. We constructed the five different Runx1/2 protein chimeras guided by AlphaFold-predicted structure to preserve domain integrity (Fig. 5D, Supplemental Fig. S4). We then transfected 200ng of each of the five plasmids into *Runx1*KO cells, and selected polyclones of transfected cells with Puromycin. When the Puromycin-resistant cells reached 80% confluence we switched them to DM for 6 days. Substituting the highly conserved N-terminal DNA-binding Runt domain (#1, Runx1^RUNT2^) resulted in small reduction of rescue activity, as did the swap of the C-terminal AD1, ID1 domain and VWRPY terminal sequence (#2, Runx1^AD/ID2^, assessed by fusion index (Fig. 5E), fiber length (Fig. 5F), and MHC and Hoechst staining (Fig. 5G)). The Runx1 RUNT domain did not rescue muscle differentiation when placed into Runx2 (#3, Runx^1/2^, Fig, 5E-G). Inserting the Runx2 MID2 domain into Runx1 (#4, Runx1^MID2^) abrogated rescue activity (Fig. 5E-G). Importantly, substituting the MID1 domain into Runx2 (#5, Runx2^MID1^) rescued fusion (Fig. 5E-G). These data indicate that the MID1 domain of Runx1 is required and sufficient to support muscle differentiation.

### RUNX1 MID interaction with ETS family member ETV4 is key to Runx1 unique functions

Our experimental results and the predicted protein structures for RUNX1, RUNX2, and the four of the RUNX 1/2 chimeras we constructed, identified a distinguishing helix present in the MID domains of the RUNX1 and absent from RUNX2 proteins. This helix in Runx1 constitutes an ETS1 interaction domain required to induce target gene expression. RUNX1/ETS1 complex has been crystalized on a composite motif target wherein the ETS motif is followed by the RUNX motif (Shrivastava et al., 2014). It has been previously shown that ETV4 (PEA3), an ETS domain-containing transcription factor, accelerates myogenic differentiation when overexpressed in vitro (Taylor et al., 1997). We hypothesized that RUNX1 MID1 domain impacts myogenesis by partnering with ETV4, thereby enhancing Runx1 relatively weak intrinsic transactivation. To test this possibility, we co-expressed a Flag-ETV4 and Myc-tagged Runx1, Runx2 and Runx1^MID2^ in C2C12 cells and performed co-immunoprecipitation experiments using magnetic beads coated with antibodies against FLAG or MYC tags. Only Runx1 protein containing the MID1 domain immunoprecipitated Flag-ETV4 (Fig. 6A).

**Figure 6:**
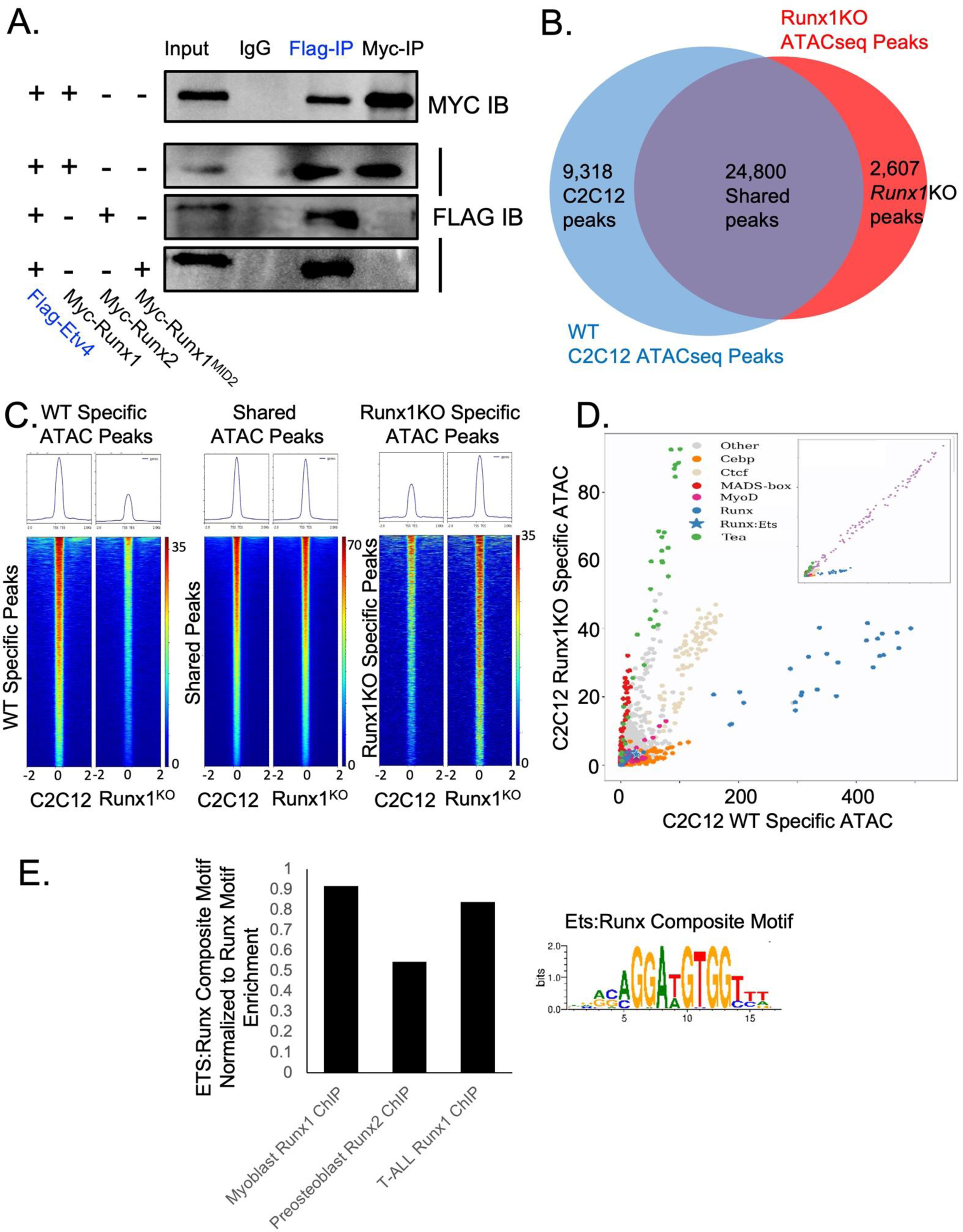
Runx1 maintains chromatin accessibility through binding Etv4. (A) Western blots from co-immunoprecipitation experiments in C2C12 cells transfected with Myc tagged Runx proteins and Flag-tagged Etv4. (B) Venn diagram of ATACseq peaks from C2C12 or Runx1KO cells. Most peaks are shared, 9,318 peaks that are only found to C2C12 and 2,607 peaks are only found in Runx1KO C2C12 cells. (C) Heatmaps showing ATACseq reads from WT or Runx1KO C2C12 cells mapped onto these 3 classes of ATACseq peaks (WT specific, Shared, and Runx1KO specific). (D) Graph displaying the -log pValue of transcription factor motif enrichment as determined by HOMER. Runx1 motifs (blue) are more highly enriched in WT C2C12 cells, whereas Mef2 motifs (red MADS-box) are more enriched in Runx1KO C2C12. The inset shows that AP1 motifs (purple) are similarly enriched in both WT and C2C12 cells as expected for a global enhancer binding factor. (E) Bar graph showing enrichment for the Ets:Runx composite motif in Runx1 ChIP data from myoblasts or T-ALL cells, less enrichment seen in Runx2 ChIP from preosteoblasts. The composite motif enrichment in each dataset is normalized to enrichment rate of the Runx-only motif in the respective ChIP from each cell types.

In order to further understand the underlying causes of the transcriptional changes observed in the Runx1KO C2C12, we performed ATAC-seq on triplicate samples of WT and Runx1KO C2C12 grown in GM and DM. Analysis of the ATAC data (https://www.ncbi.nlm.nih.gov/geo/query/acc.cgi?acc=GSE248044) identified 24,800 peaks that were consistent in all replicates and shared between control and Runx1KO C2C12 cells (Fig. 6B). Importantly, 9,318 ATAC-seq peaks present in all C2C12 triplicates were not found in Runx1KO cells. Conversely, 2,607 ATAC-seq peaks were unique to Runx1KO (Fig. 6B). This is demonstrated visually by mapping all ATAC-seq reads from C2C12 and Runx1KO cells to either shared or unique peaks sh (Fig. 6C). The higher number of ATAC-seq peaks unique to C2C12 and lost in Runx1KO cells points again to Runx1 functioning primarily as a transcriptional activator. Motif enrichment analysis with the software package HOMER showed that the Runx1 motif is highly enriched in C2C12 specific peaks consistent with a role for Runx1 in keeping these chromatin regions open (Fig. 6D, (Hass et al., 2021)). Genomic snapshots of the *Col6a1*, *Col6a2*, and *Cdh15* display reduced chromatin accessibility in the Runx1KO C2C12 cells compared to WT C2C12 cells, in the same regions that had reduced accessibility in Runx1KO mK4 cells (Fig. 2H and I).

To ask whether the interaction between ETV4 and Runx1 contribute to chromatin regulation by in C2C12, we tested for presence of the Ets:Runx composite motif utilizing the MCOT anchored motif analysis that searches ATAC-seq peaks for any secondary motifs at variable distance and orientation from a Runx motif. The MCOT program identified significant enrichment of an Ets motif adjacent to a Runx1 motif in C2C12 unique peaks with orientation and spacing identical to the known Ets:Runx composite motif (p-value 1 x 10^−14^) (Supplemental Fig. S6A). The Ets:Runx motif enrichment in the Runx1KO unique peaks was negligible. The enrichment of the Ets:Runx composite motif in the C2C12 unique peaks was confirmed using the MEME Suite Simple Enrichment Analysis (SEA) tool (p-value 4.37 x 10^−22^). This suggests that the Runx1 and ETV4 may be cooperating to keep chromatin open and drive expression of a subset of targets genes. Among the Runx1KO unique peaks, one of the most enriched motif was the *Mef2c* motif (Fig. 6D), consistent with the observation that *Mef2c* is upregulated in the Runx1KO cells and suggesting that even the low level of Mef2c expression in GM is sufficient to alter accessibility to chromatin regions in the absence of *Runx1*.

To ask if any C2C12-unique ATAC-seq peaks are directly regulated by Runx1 binding, we intersected our C2C12- and the Runx1KO-unique peaks with a published Runx1 ChIP from myoblasts (Umansky et al., 2015). We found that 3938 of the 9318 (42.3%) of the C2C12 unique peaks were bound by Runx1 in myoblasts, while only 21.2% (552 out of 2607) of Runx1KO unique ATAC-seq peaks were bound by Runx1 (Supplemental Fig. S5A and B). Consistent with our previous report (Hass et al., 2021), direct binding by Runx1 appears to maintain chromatin accessibility at many Runx1 and Ets:Runx specific ATAC-seq peaks.

To explore whether combinatorial binding of ETS and Runx1 to the composite Ets:Runx site could underlie the unique functions of Runx1 in myoblasts, we tested the prediction that the composite Ets:Runx site will represent a larger fraction of all RUNX sites in cell types that depend on *Runx1* but not in a cell type that is dependent on *Runx2* in three ChIP-seq data sets showing highly significant enrichment of Runx motifs. In a Runx1 ChIP-seq dataset generated in myoblasts, composite motif enrichment near genes related to muscle development is highly significant (p-value 2.11 x 10^−9^) (Supplemental Fig. S5C). Likewise, in a Runx1 ChIP-seq datasets generated from the T-ALL cell line CUTLL1, very significant enrichment (p-value 2.42 x 10^−21^) for the composite motif was observed near genes regulated during T-cell development. By contrast, composite motif enrichment near genes regulating bone development was modest (p-value 6.77 x 10^−2^) in Runx2 ChIP-seq datasets generated from preosteoblasts. Importantly, all three ChIP data sets showed highly significant enrichment of Runx motifs (myoblast Runx1 ChIP, *p*=4.74×10^−57^; T-ALL Runx1 ChIP *p*=1.95×10^−28^; preosteoblast Runx2 ChIP *p*=2.61×10^−32^), highlighting the enrichment specificity of the composite motif in Runx1 ChIP (Fig. 5E and Supplementary Fig. S5D). Taken together, these analyses support the idea that *Runx1*-dependent tissues rely on the combinatorial function of Ets and Runx1 proteins.

## Discussion

Runx1 had previously been shown to be required for myoblasts differentiation during muscle regeneration from injury, but the role of Runx1 in quiescent SCs was not addressed during development or normal muscle maintenance. Deleting *Runx1* in SC using an inducible Pax7-Cre in uninjured muscle demonstrated that Runx1 is required to maintain SC throughout life, in part by keeping Notch-depended enhancers accessible. Runx1 is thus required for all phases of skeletal muscle development, maintenance and regeneration.

Analysis of Runx1KO C2C12 revealed that the muscle transcription factor Mef2c was upregulated in growth media with dramatic upregulation during differentiation, noted also during differentiation of Runx1KO myoblast (Umansky et al., 2015). Increasing Mef2c levels artificially in in WT C2C12 generated a similar fusion blockade, which suggests that loss of fusion in Runx1KO c2c12 cells may be caused by the inappropriate upregulation of Mef2c. It is possible that Runx1 directly represses *Mef2c* expression; alternatively, as we recently reported (Hass et al., 2021) much of the repressive role of Runx1 in mK4 is indirect, mediated through the Runx1 targets *Zeb1/Zeb2*. We noticed that *Snai1,* a transcriptional repressor of *Mef2c* in C2C12 (Batlle et al., 2013), is downregulated in C2C12 and may be direct Runx1 target. This raises the possibility that the increased Mef2c levels and activity in Runx1KO is an indirect outcome of losing *Runx1* and its target, *Snai1*.

Despite high degree of conservation and affinity to highly related binding motifs, co-expressed Runx proteins have demonstrable non-redundant functions. *Runx1* has been extensively investigated for its role in T cell development, where it is co-expressed with *Runx3*, and in muscle development, where it is co-expressed with *Runx2*. Similar to its multiple roles supporting the common lymphocyte progenitors through their subsequent differentiation to T and B cell linages in adult hematopoiesis (Growney et al., 2005; Rothenberg, 2021; Shin et al., 2021), we show here that in addition of an established role in the adult skeletal muscle response to injury, *Runx1* is also involved in SC maintenance in the absence of injury.

Neither one of these functions are satisfied by the resident Runx2. Deletion of *Runx1* specifically in the SC reduced their numbers in uninjured adult mice (defect in self renewal) and impaired muscle regeneration (failure of myoblasts to differentiate). In part, the role of Runx1 in myoblasts involved a cooperative function with MyoD. However, *MyoD* is not expressed in quiescent satellite cells suggesting that Runx1 functions in SC depend on other factors with whom Runx2 cannot interact. In T-cells, Runx1 interacts with Ets family transcription factors, and accordingly, a composite Ets:Runx motif was highly enriched (*p=*10^−21^) in Runx1 ChIP-seq datasets generated from T-ALL cells. We observed that Runx1*-* dependent ATAC-seq peaks present in C2C12 myoblast but absent from Runx1KO were also strongly enriched (*p=*10^−22^) for the composite motif, with similar enrichment detected in Pax7 ChIP-seq datasets from ES cells undergoing muscle differentiation, and in Runx1 ChIP-seq datasets generated from myoblasts (*p=*10^−9^). The enrichment of the composite Ets:Runx motif in Runx2 ChIP-seq datasets generated from preosteoblasts was orders of magnitude weaker (*p=*10^−2^).

Runx/ETS interactions observed in crystal structure of the Runx1 and Ets1 on DNA containing the composite motif are mediated by helix within a conserved region termed MID. AlphaFold predicted an alpha helix in the Runx1 MID (MID1), but not in Runx2 MID (MID2). The binding of Runx1 to Ets1 prevents Ets1 autoinhibition leading to enhanced targets transcription (Shrivastava et al., 2014). We found that the MID1 domain was both necessary for Runx1 and sufficient when inserted into Runx2 to rescue myoblast differentiation/fusion defect in *Runx1*-deficient C2C12 cells. Furthermore, we found that Runx1, but not Runx2, is able to co-immunoprecipitate Etv4, an Ets family member that has previously been implicated in myoblast differentiation. Similarly, in mK4 cells, both *Runx1* and *Runx2* are expressed; yet *Runx1* deficiency caused dramatic changes in chromatin accessibility despite Runx2 binding to many of the regions that closed in the absence of *Runx1* (Hass et al., 2021).

Runx proteins collaborate with many TFs at different stages of development, most famous among them is the CBFβ transcription factor binding to Runx proteins through the proline-rich (PY) repeat. We propose that non-redundant activity of *Runx1* in satellite cells, T-cells, and perhaps additional tissues or developmental stages reflects its ability to form a sable complex with Ets transcription factors via the MID domain. This complex can maintain chromatin accessibility and facilitate transcriptional regulation of essential targets genes via the Ets:Runx composite motif recruiting additional coactivators (Hollenhorst et al., 2009). However, the role of the Ets:Runx composite sites is rather complex: the Ets protein *Fli1* blocks effector T-cell differentiation (T_EFF_) by occluding Ets:Runx composite sites. loss of *Fli1* enables T_EFF_ differentiation in a *Runx3* dependent fashion (Chen et al., 2021). *Runx3* is not predicted to form a complex with Ets by AlphaFold, perhaps binding to this site without an Ets partner. By contrast, *Runx1* over expression in Fli1-defiecint cells reverses the gains in T_EFF_, promoting an alternative T-cell type (Chen et al., 2021), perhaps by recruiting another Ets protein to the composite site. It will be interesting to see if (1) *Runx2* can rescue a *Fli1, Runx3* deficient T-cell in the experimental paradigm explored by Chen et al, and if Runx1Mid2 chimera will promote T_EFF_, like *Runx3*, instead of suppressing them.

## Materials and Methods

### Mice

All mice used in this study were on the C57BL/6J/CDI mixed genetic background, between 8– 12 weeks, and had age-matched littermate controls. Both sexes were used. Mice maintained at Cincinnati Children’s Hospital Medical Center animal facility were handled following animal care guidelines in a were approved by the Animal Studies Committee of CCHMC (IACUC 2018-0108/0107). After transfer to Fudan University mice were handled following animal care guidelines approved by the Fudan Animal Studies Committee (IACUC IDM20211021). *Runx1*^flox/flox^ mice were crossed with *Pax7*^CreERT2/+^*; Runx1^+^*^/flox^ mice (Lepper et al., 2009) to generate *Pax7^CreERT2/+^; Runx^1fl/fl^*mice. Tamoxifen (Aladdin) was dissolved in 90% (vol/vol) sesame oil and 10% (vol/vol) ethanol at 20 mg/mL and delivered to both control and Runx1cKO mice at 8 weeks of age by i.p. injection at 0.1 mg/g every day for 5 days. Genotyping primers are in Supplemental Data 1 file.

### TA muscle Cardiotoxin Injury

Cardiotoxin (CTX) (Latoxan) was dissolved in sterile saline solution to a final concentration of 10 μM, aliquoted, and stored at −80 °C. For muscle injury, mice were anesthetized by i.p. injection of 2.5% Avertin (Aladdin) at (15 μL/g). The fur on the hind legs was shaved and the legs were swabbed with 75% ETOH. Intramuscular injections into the TA muscle of 50 μL of CTX were performed with a 26-gauge needle. As a control, contralateral TA muscles were injected with the same volume of normal saline. After injection, animals were kept under a warming lamp until the anesthetics wore off. Post injection mice were anaesthetized and euthanized by CO2 asphyxiation to harvest the TA muscles at 5, 7, 12 and 30 days post-injury. Tissue was fixed in 4% PFA for 12-18h, dehydrated in ethanol/xylene series and embedded in Paraffin. Paraffin sections were rehydrated for histological analyses as detailed below.

### Hematoxylin and eosin (H&E) staining

For the assessment of muscle morphology, 5 μm-thick cross-sections of TA muscles were subjected to H&E staining which was performed according to standard procedures provided by the H&E staining kit (Servicebio).

### C2C12 cell culture

C2C12 myoblasts were purchased from ATCC and proliferated in Dulbecco’s Modification of Eagle’s Medium (Gibco, DMEM) with 1% Pen/Strep (Gibco) and 20% (v/v) fetal bovine serum (Sigma, growth medium, GM) under moist air with 5% CO_2_ at 37 °C. Post-confluent C2C12 cells were washed with DPBS (Gibco) and incubated with DMEM supplemented with 1% Pen/Strep and 2% (v/v) horse serum (Gibco, differentiation medium, DM) at the same condition for differentiation. Inducing cell differentiation was performed when cells were confluent to make sure the same cell density.

### Generation of Runx1KO C2C12 cells

We cloned guide RNAs (sgRNAs, Supplemental Data 1) targeting Exon 3 of *Runx1* that contains the start codon in the px459 CRISPR/Cas9 vectors. The sgRNA plasmids were constructed with the method and tools described in (Haeussler et al., 2016; Ran et al., 2013). We split C2C12 cells into control and experimental sibs and transiently transfected pX459-sgRNA plasmids into the experimental sib with Lipofectamine 2000 (Invitrogen) according to the manufacturer’s instructions. At 12 hours post-transfection, the cells underwent selection with 5ug/mL puromycin for two days. When all cells alive in the control dish died, clones growing on the transfected were picked ∼ 1 week later using cloning disks. The clones were tested for Exon3 deletion by the T7E1 assay and for Runx1 protein expression by Western blot. These cells were characterized in Cincinnati for the data shown in Figure 3; however, *Runx1*KO-C2C12 were independently recreated by the same investigator (MY) in Fudan University using the same protocol with cells from a different stock of C2C12. The results presented in Figure 4-6 were reproduced at both institutions, the molecular analysis was performed on the Fudan *Runx1*KO-C2C12 cells.

### Western blot

Confluent C2C12 control or *Runx1*KO cells were collected in 100 ul of RIPA lysis buffer (Beyotime) with protease inhibitors (Bimake) and PMSF (LIFE-iLAB) plus 20 ul of 6X sample buffer. Protein samples were run on EZ Protein any KD PAGE (LIFE-iLAB) and then transferred to 0.22um PVDF membranes (Millipore). Membranes were blocked using 5% non-fat milk in PBS-Tween20 0.1% for 1 h at room temperature. Indicated primary antibodies (Supplemental Data 1) were applied at 1:1000 dilutions overnight at 4 degrees, and then HRP-conjugated secondary antibodies were used at 1:2000 at room temperature for 1hr. The Western blot signal was detected using Super ECL Detection Reagent (Yeasen) using a Bio-Rad Chemidoc MP Imaging System.

### RNA extraction and Quantitative real-time PCR

RNA from biological triplicate samples were extracted from C2C12 cells with PuroLink kit (Invitrogen) and the Reverse transcription kit (Promega) was used to make cDNA following manufacturer indications. The cDNA was diluted to 20 ng/ul and 2ul of each sample was added to each qRT-PCR reaction that was amplified using 2x SYBR Green qPCR Master Mix (Bimake) and performed on a CFX96^TM^ Connect Real-Time PCR System from Bio-Rad. Gene expression levels were normalized to mouse *Gapdh* expression and changes were determined relative to control cells with significance calculated using the two-way Student’s t-test.

### Primary myoblast purification and culture

Isolation of primary myoblasts was performed as described before (Hindi et al., 2017). Briefly, hind limb muscles were pooled, minced, and digested with 400U/mL Collagenase II (Yeasen), followed by trituration. Cell suspension was filtered through 70- and 40-μm cell strainers (Biosharp), and primary myoblasts were pelleted after centrifugation. Isolated primary myoblasts were cultured on 10% Matrigel (Corning) -coated 10cm dishes. Proliferation medium contained Ham’s-F10 (Gibco) with 1%Pen/Strep, 20%FBS (Myoblasts growth medium) and 2ng/mL basic human fibroblast growth factors (FGF2). When cell confluency has reached ∼70%, the cells media was replaced with differentiation medium (DMEM with 2% horse serum and 1%Pen/Strep).

### Immunofluorescence staining

Cells were cultured in Chamber Slides (Thermo Fisher) or 12-well plates and were fixed in 4% paraformaldehyde (PFA) for 15min and permeabilized in 0.3% PBS-Triton X−100 for 3X10min at room temperature. Isolated TA muscles were fixed in 4% PFA at 4°C overnight, dehydrated by graded ethanol and embedded in paraffin. Paraffin-embedded samples were cut into 5-μm sections using Leica RM2125RTS. Samples were then blocked in 10% Normal Donkey Serum (Jackson ImmunoResearch Laboratories) in PBS for 1h at room temperature followed by incubation with primary antibodies at 4°C overnight. Subsequently, appropriate fluorescently labeled secondary antibodies (Alexa Fluor 488, or 549) were incubated for 1 h at room temperature. Hoechst was used to stain the cell nuclei for 15 min. Images were acquired with confocal microscope (Olympus). Antibodies are listed in Supplemental Data1 file.

### Transfection of plasmids

The mouse *Runx1* overexpression plasmid pCDNA3.1-Flag-Runx1 was purchased from Addgene. The mouse Runx2 plasmid pCMV-Flag-mRunx2 was purchased from Origene. The fragments shown in Fig. 5D were cloned from pCDNA3.1-Flag-Runx1 or pCMV-Flag-mRunx2 by using Phanta Super-Fidelity DNA Polymerase (Vazyme) with primers are listed in Supplemental Data 1. ClonExpress II One Step Cloning Kit (Vazyme) was used to ligate and construct RUNX1/2 chimeras. Sequences were confirmed by Sanger sequencing performed by Beijing Tsingke Biotech Co., Ltd. For the rescue experiments and chimeras transfection, 200ng plasmids (a mix containing the Runx plasmid and pLVX-GFP) were transiently transfected into 12-well plates according to the protocols recommended by Lipofectamine 2000 manufacturer (Invitrogen). For Mef2c overexpression experiment, confluent cells were transfected twice with a 24 h interval using Lipofectamine 3000 (Invitrogen) according to the manufacturer’s instruction. pCDNA3.1 was used as an empty vector transfection control. Transfection efficiency was monitored using GFP fluorescence.

### Fusion Index, Quantification and Myotube dimensions Analyses

C2C12 cells and isolated primary myoblasts were cultured in the differentiation medium to induce terminal differentiation. Histological parameters were derived from on at least four different animals per genotype. Phase-contrast and immunofluorescent images were obtained in four randomly selected fields in each of three biological replicates using a 20× objective and then analyzed using ImageJ software.

### RNA-seq generation and analysis

C2C12 were grown and differentiated as described above. RNA was isolated by using Invitrogen’s Purelink RNA Mini kit according to manufactures directions. The DNBSEQ Transcriptome libraries were performed by BGI Hongkong Tech Solution NGS Lab to produce over 20 million reads per sample. Raw RNA-seq data was processed using a pipeline developed in-house called CSBB (https://github.com/csbbcompbio/CSBB-v3.0). It employs fastqc to check read quality followed by Bowtie2 + RSEM for alignment and quantification against the mm10 genome. The final outputs are TPM and counts matrices at the gene and isoform level. Differentially expressed genes were then identified using DESeq2 (doi:10.1186/s13059-014-0550-8).

### ATAC-seq data generation and analysis

C2C12 were grown and differentiated as described above until nearly confluent. Cells in GM were washed and media replaced with lysis buffer (10 mM Tris-HCl (pH 7.4), 10 mM NaCl, 3 mM MgCl2, 0.5% NP-40) for 10 min on ice, then spun at 500 g for 5 min at 4℃ to obtain nuclei. Nuclei were then segmented with Tn5 transposase (Vazyme) at 37℃ for 30 min. The ATAC libraries were prepared by using TruePrep DNA Library Prep Kit V2 for Illumina (Vazyme, TD501) according to manufactures direction. VAHTS DNA Clean Beads (Vayzme, N411) was used to do the libraries selection. An aliquot was tested by Agilent 2100 Bioanalyzer for average fragment size. Libraries were sequenced on Illumina Novaseq 6000 by NJNA Biopharmaceutical Public Service Platform.

Paired reads were processed with the nf-core ATAC-seq pipeline v2.0 (<u>10.5281/zenodo.2634132</u>), using the mm10 genome/blacklist and specifying parameters “– read-length = 100” and “–narrow-peak”. It encompasses all steps from FASTQ trimming and quality control to final bigwig and peak files. The main tools used include BWA for alignment, MACS2 for peak calling, and HOMER for annotation. To identify differentially accessible peaks, the individual ATAC-seq peak files were intersected using the Bedtools intersect intervals tool on https://usegalaxy.org to identify peaks that were present in all triplicates. The same tool was then utilized to identify the shared peaks between WT and Runx1KO cells and the peaks unique to each. The peaks from the different classes were analyzed using HOMER to determine enrichment of transcription factor motifs. The heatmaps of the ATAC-seq reads were generated using the Deeptools computeMatrix and plotHeatmaps tools on https://usegalaxy.org. GEO the peak bed files for myoblast Runx1 ChIP (GSE56077), preosteoblast Runx2 ChIP (GSE54013), and CUTLL1 Runx1 ChIP (GSE51800) were annotated to nearest genes using the Great Annotation tool and intersected with the gene lists from GO terms “muscle differentiation”, “bone development”, or “T cell differentiation”, respectively. The Runx1 or Runx2 ChIP peaks were subjected to motif enrichment using HOMER and MEME-Suite SEA enrichment tools to determine enrichment of Ets:Runx composite site, or submitted to the MCOT program and searched for motifs near the Runx1 anchor.

### Statistics

Data are presented as mean ± SEM. Differences between groups were tested for statistical significance by using the unpaired two-tailed Student t-test (Excel& GraphPad Prism Software). P < 0.05 was considered significant. P<0.05 was marked with an asterisk (*), P<0.01 was marked with double asterisks (**) and P<0.001 was marked with triple asterisks (***). For multiple comparison analysis, FDR<0.05 was selected. Analysis-specific cutoffs are described in the supplemental table presenting the data.

### Data Availability Statement

In addition to source data presented in Supplemental Data Table, RNA and ATAC data are deposited here https://www.ncbi.nlm.nih.gov/geo/query/acc.cgi?acc=GSE248046.

## Acknowledgments

The authors wish to thank Dr. Doug Millay and members of the Kopan and Lin laboratories for constructive discussions.

## Authors contributions

Study conception and design: RK and MH. Experimental plan: MH, RK, MY. Data collection: MY, WW, CY. Data analysis and interpretation of results: RK, MH, MY, XL; Draft manuscript preparation: MY, RK. and MH; Bioinformatic analyses: MH, SP and KT. Project supervision in Fudan; XL. All authors reviewed the results and approved the final version of the manuscript.

