## Supplemental Data 1 for "The unique function of Runx1 in skeletal muscle differentiation and regeneration is mediated by an ETS interaction domain"

**Supplementary Data 1**

Table S1

**PCR and sgRNA Primers.**

| Runx1 exon3 gRNA F1 | CACCGGCTCCTACTAGACGGCGAC |
| --- | --- |
| Runx1 exon3 gRNA R1 | AAAC GTCGCCGTCTAGTAGGAGCC |
| Runx1 exon3 gRNA F2 | CACC GCGGTGCGCACTAGCTCGCC |
| Runx1 exon3 gRNA R2 | AAAC GGCGAGCTAGTGCGCACCGC |
| Runx1 exon3 genotyping F | GAGGTGAGAGAGTTGACCTGGAAAC |
| Runx1 exon3 genotyping R | GGTGAACCCTCTGTGTGCATTACA |
| Runx1 flox genotyping F | GAGTCCCAGCTGTCAATTCC |
| Runx1 flox genotyping R | GGTGATGGTCAGAGTGAAGC |

**RT-qPCR primers**

| myogenin-F | CTACAGGCCTTGCTCAGCTC |
| --- | --- |
| myogenin-R | ACGATGGACGTAAGGGAGTG |
| MEF2C-F | GCCGGACAAACTCAGACATT |
| MEF2C-R | TGGGATGGTAACTGGCATCT |
| GAPDH-F | CATCACTGCCACCCAGAAGACTG |
| GAPDH-R | ATGCCAGTGAGCTTCCCGTTCAG |
| MyoD-F | TACAGTGGCGACTCAGATGC |
| MyoD-R | GAGATGCGCTCCACTATGCT |
| Myosin-F | CGTTTTGGACATTGCGGGTT |
| Myosin-R | ACCGTCCGCATCTGTTGTAG |
