## Supplemental Figures for "The unique function of Runx1 in skeletal muscle differentiation and regeneration is mediated by an ETS interaction domain"

**Supplemental Figure S1:**

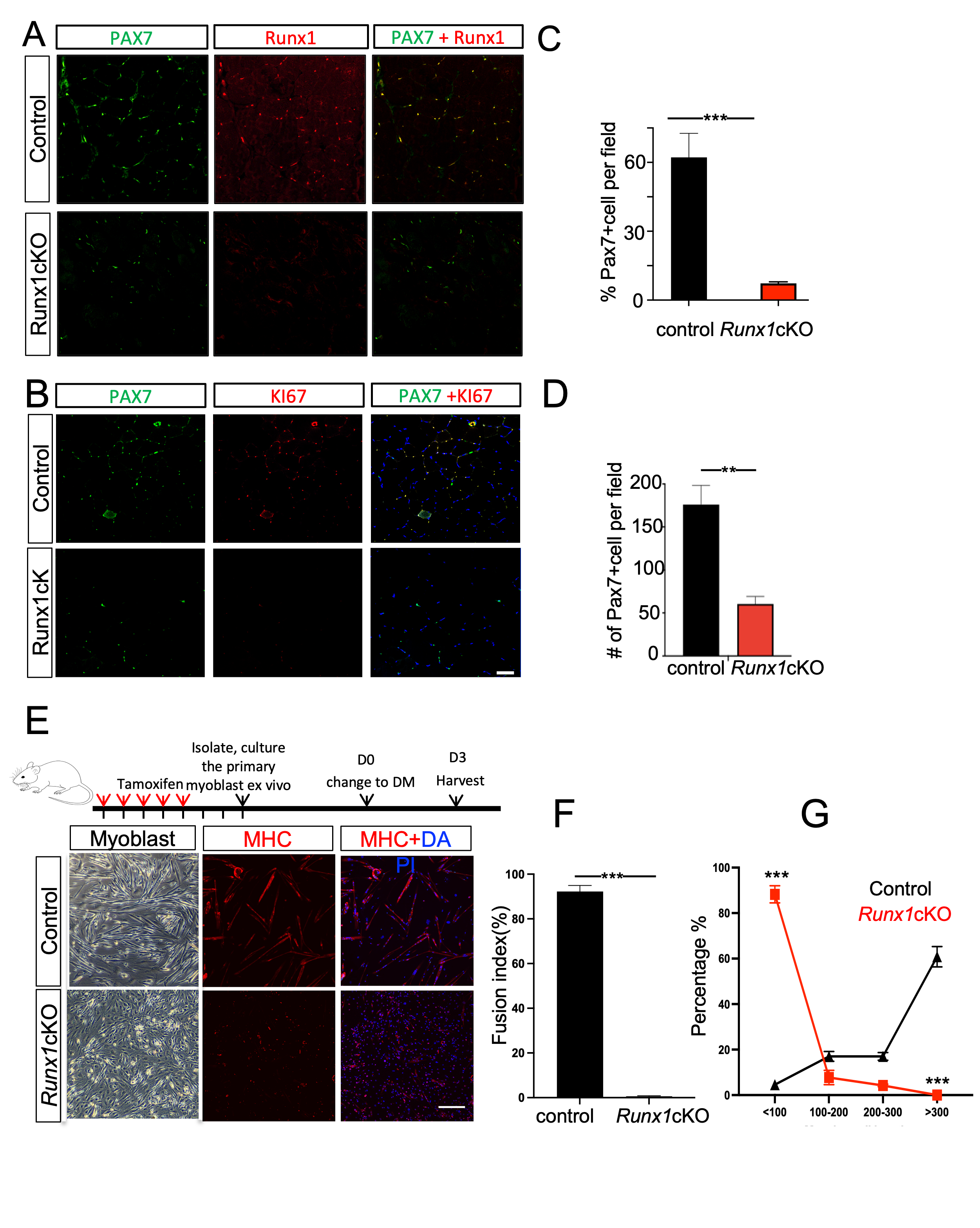

Supplemental Figure S1: (A) Immunostaining of Pax7 and Runx1 in TA muscle cross-sections after 5 consecutive daily tamoxifen injections. (B) Immunostaining of Pax7, Ki67, and DAPI (blue) in TA muscle cross-sections on day 5 post-injury. (C) Quantification of Pax7-positive cells per field in cross sections of TA muscle from control and Runx1cKO mice as shown in (A). (D) Quantification of Pax7-positive cells per field in cross sections of TA muscle from control and Runx1cKO mice as shown in (B). (E) Representative phase contrast microscopy images of isolated primary myoblast from control and Runx1cKO mice cultured for 7 d in proliferation medium followed by 3 d in differentiation medium. Representative images of immunofluorescence staining for Myosin heavy chain (MHC) in differentiated, isolated primary myoblasts. Nuclei were stained with DAPI. Quantification of fusion index (F) and MHC+ cell lengths (G) in experiments shown in E. Data are represented as mean ± SD. n=4 for each group. **P < 0.01; ***P < 0.001 (Student’s t-test). Scale bars: 80μm.

**Supplemental Figure S2:**

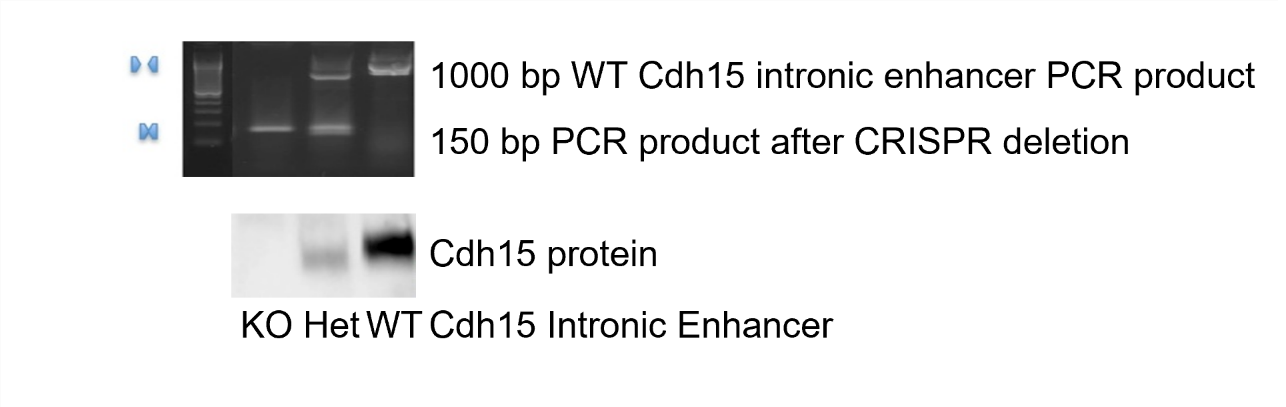

Supplemental Figure S2: Top panel shows PCR spanning the Cdh15 intronic enhancer showing the expected 1000bp size in WT mK4 cells or 150 bp band expected after the CRISPR mediated deletion in the KO cells, while heterozygous cells had both bands. Bottom panel is the Western blot for Cdh15 in mK4 cells with Notch signaling activation inducing expression of Cdh15 in WT cells, only partially in heterozygous cells and a complete lack of expression of Cdh15 in cells where the intronic enhancer has been deleted.

**Supplemental Figure S3:**

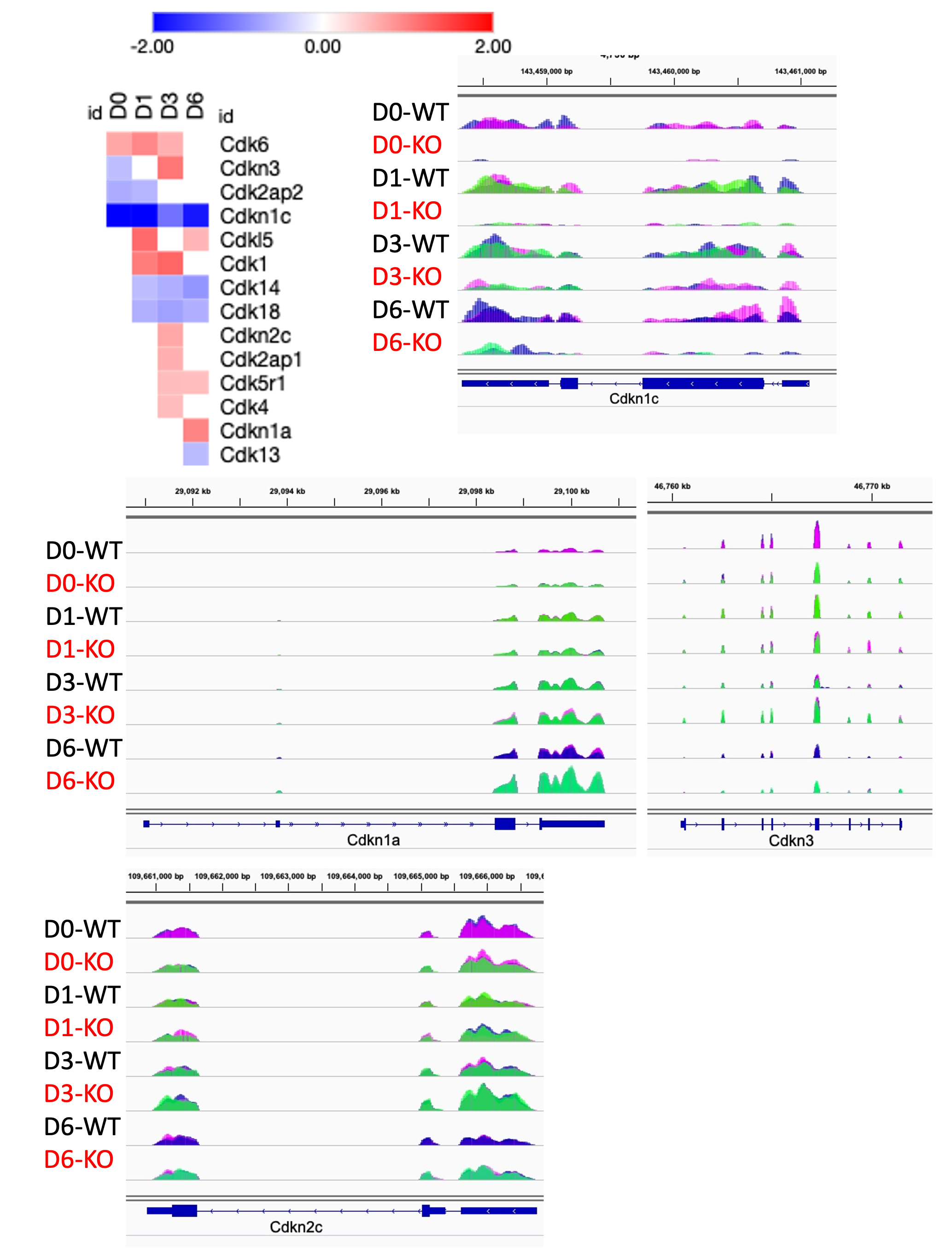

Supplemental Figure S3: Heatmap and genomic snapshots of RNA reads for indicated genes from WT and Runx1KO C2C12 cells grown in growth media or for different times of differentiation media.

**Supplemental Figure S4:**

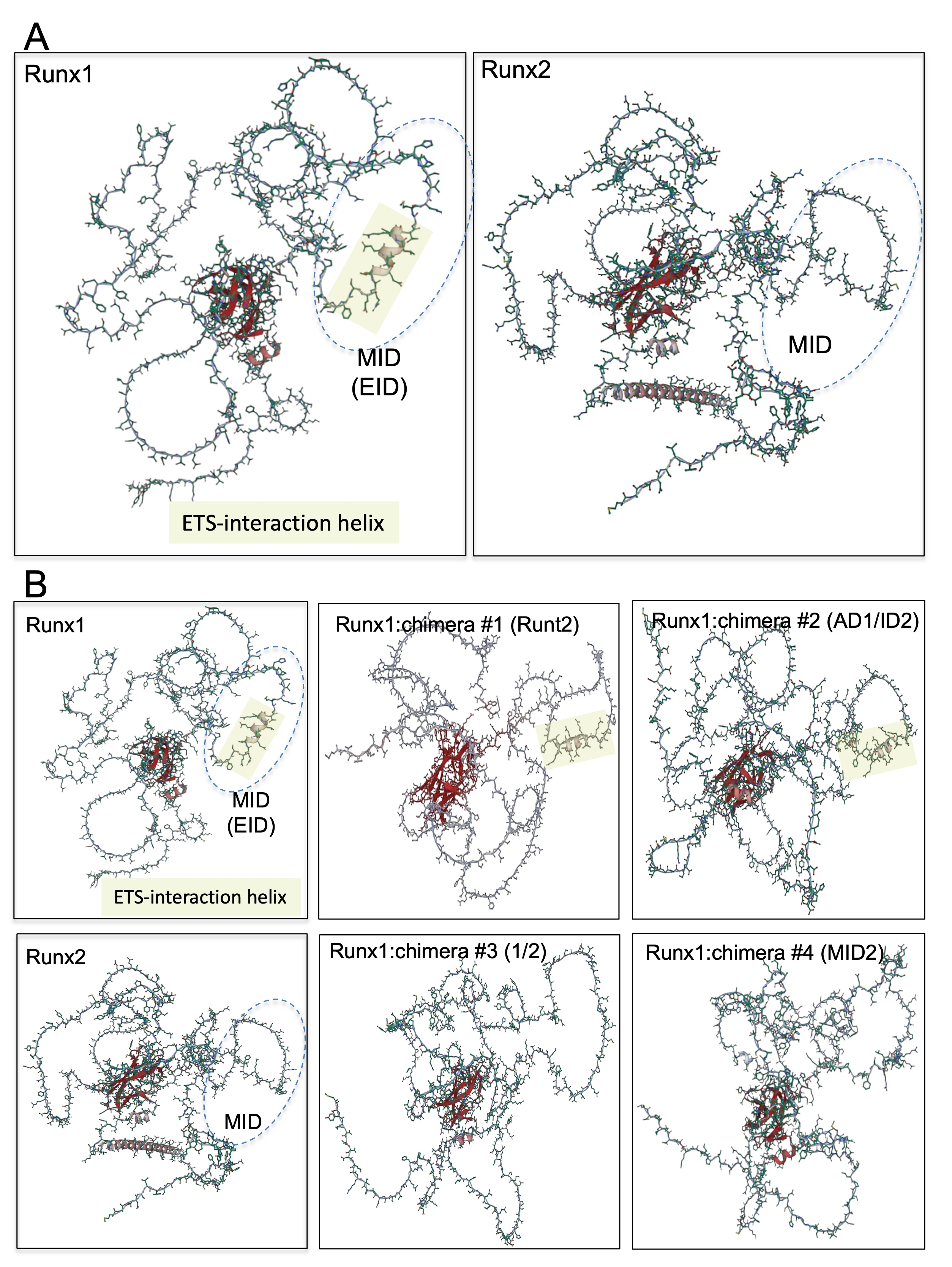

Supplemental Figure S4: (A) AlphaFold images of Runx1 or Runx2 protein structure showing the ETS1-binding alpha helix in Runx1 (shaded). (B) AlphaFold images of Runx1, Runx2, and the various Runx1/2 chimera proteins used in rescue experiments. The ETS interaction domain is shaded.

Supplemental Figure S5:

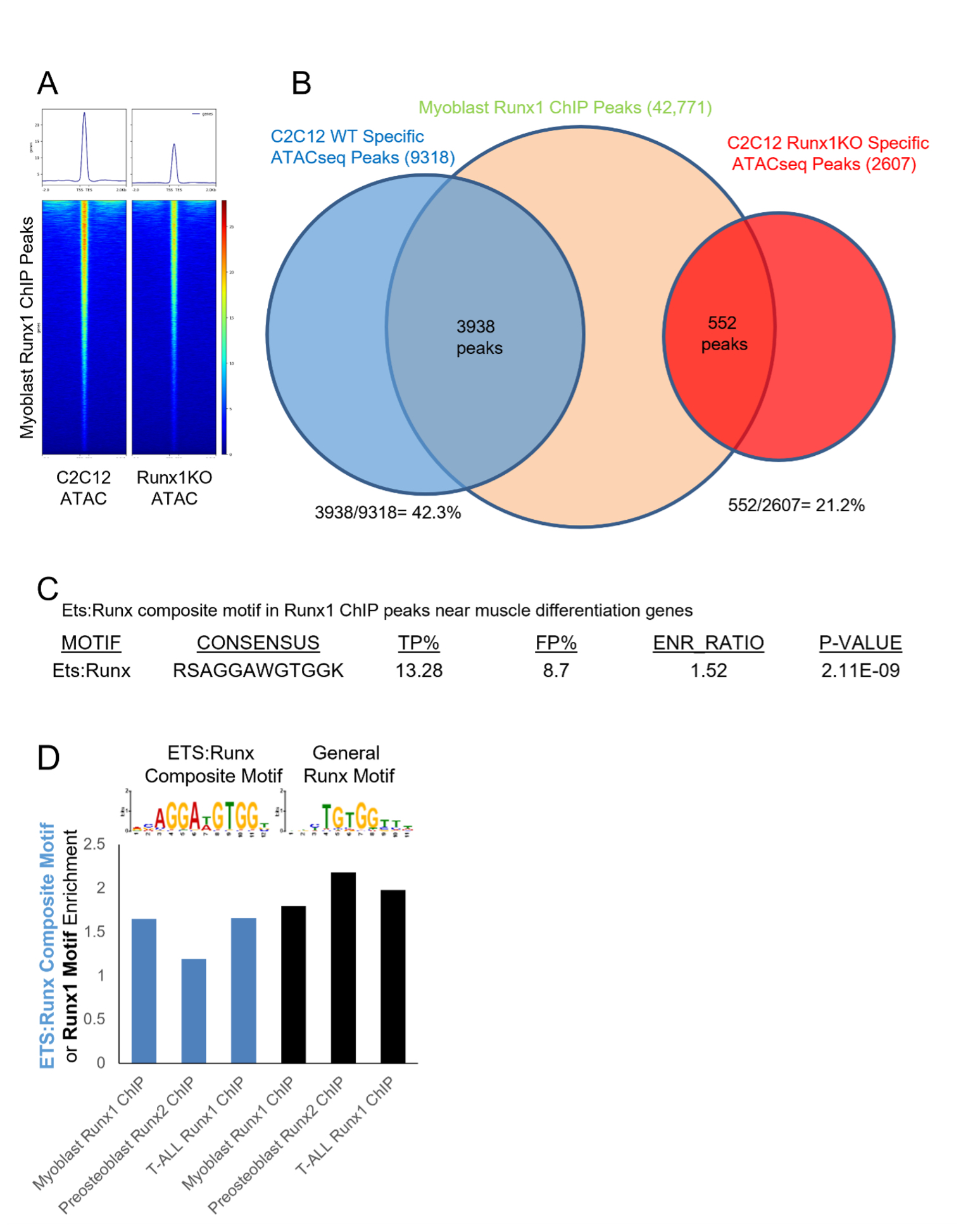

Supplemental Figure S5: (A) Heatmaps displaying ATAC-seq reads from control or Runx1KO C2C12 cells mapped onto Runx1 ChIP peaks from myoblasts. (B) Venn diagram showing the overlap of myoblasts Runx1 ChIP peaks with ATAC-seq open chromatin regions from control C2C12 cells (42.3% overlap) or from Runx1KO C2C12 cells (21.2% overlap). (C) Simple Enrichment Analysis (SEA) determined that Ets:Runx composite motif was significantly enriched in myoblast Runx1 ChIP near muscle differentiation genes. (D) Bar graph showing the enrichment rate of the Ets:Runx composite motif (blue) or Runx motif alone (black) that suggests that the Ets:Runx composite motif is specifically enriched in Runx1 ChIP from myoblasts or T-ALL cells but not in Runx2 ChIP from preosteoblast. The Runx motif was enriched in all the ChIP datasets. This data was used to generate the ratio shown in Fig. 6E.

**Supplemental Figure S6:**
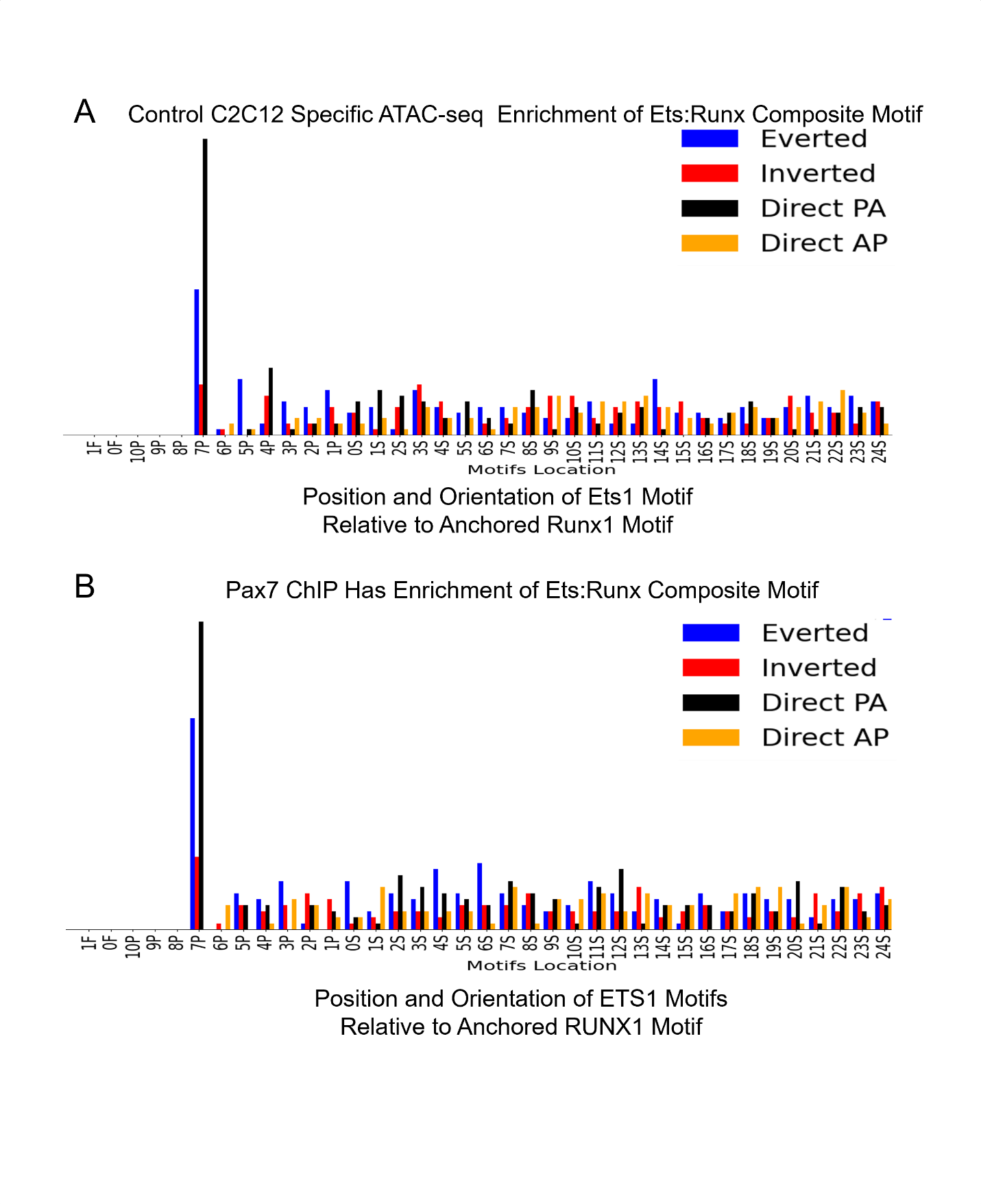

Supplemental Figure S6: (A) Motif Co-occurance Tool (MCOT) plot of the ETS1 motifs position and orientation relative to the anchored RUNX1 motif with significant over-representation of an Ets motif directly preceding the Runx motif, which conforms to the Ets:Runx composite motif. (B) MCOT plot showing enrichment of Ets motifs adjacent to Runx motifs as found in the Ets:Runx composite motif in Pax7 ChIP performed during muscle differentiation.
